# Numerical Model for Formation and Evolution of the Bleb

**DOI:** 10.1101/2020.07.06.189738

**Authors:** J. Feng, L. Tang, Z. Liu, S. Dong, L. Zhou, Y. Liu, Z. Jiang

## Abstract

The bleb morphology and its changes are an important mechanism of cell’s amoeboid migration. By releasing bonds between the membrane and the cortex of a cell, the formation of bleb can be observed experimentally, but the mechanism that affects the size and shape of this kind of bleb is waiting for further study. In this paper, a two-dimensional fluid-solid coupling model is established to describe a cell with membrane, cortex and cytoplasm in a solution, and a numerical solving method for the fluid-solid coupling model is developed to simulate the behaviors of cell bleb. The effects of parameters, such as the number of broken bonds, the viscosity coefficient of the cortex, and the cell’s membrane modulus on the size and the shape of the bleb were investigated. Numerical results show that the model is effective to simulate the formation and evolution of cell’s bleb, and derive the contribution of several affecting factors to the bleb shape and size clearly.

**SIGNIFICANCE:** To understand the process of cell migration with bleb pseudopods in the amoeba cell migration, it is necessary to study the formation mechanism of cells protruding bleb. In this paper, we propose a reasonable and reliable cell numerical model. With this model we successfully simulate the bleb phenomenon consistent with the experimental phenomenon by changing the key impact factors. The method in this paper is applicable to the cell model of amoeba cell migration pattern, which helps to understand the important role of blebs in the process of cell migration.

## 1. INTRODUCTION

Cell’s migration is a complex biochemical-physical process involving many intracellular and extracellular environmental factor (1). Generally, there are two main types of single-cell migration, namely the amoeboid migration and the mesenchymal migration (2). The amoeboid migration mode is mainly suitable for small round cells with a diameter of about 8 to 30 microns, which has a faster migration speed and consumes less energy for directional cell migration (1). In the physiological environment of the organism, different types of cells will adopt the amoeboid cell migration for deformation and migration, such as immune system cells (3), epithelial cells (4), and individual migration cancer cell (5).

The cortex is a grid of cellulose cross-arranged within the cell membrane, composed of actin fibers, myosin and related protein molecules (6–10), which exhibits strong fluidity and contractility. For amoeboid-type cell migration, the cell movement is driven by the formation and contraction of bleb. The depolymerization of the actin cortex at the front end of the cell causes the changes of the hydrostatic pressure within the cell, the pressure drives the separation of the frontal cortex from the cell membrane layer to produce a bubble-like or plate-like pseudopod, and the polymerization of the action cortex leads to contraction, causing the cells to move in a specific direction (11–12). However, the mechanism of cell bleb producing is not very clear. According to existing experimental studies, it is believed that the generation of bleb is related to cytoplasmic rheology, mechanical properties of the cortex, and cell membrane tension (6, 13–15).

Bleb plays a very important role in many biological physiological processes. It is often seen in many biological physiological processes, such as cell proliferation (16–17), cell migration (18), cell division (14, 19–20), and apoptosis (20).

At present, many mechanics studies have been performed on cells that conform to the amoeboid migration including set simplified model on the profile of bleb formation, and summarized that the force-relationship between adhesion (21–23), contraction and polymer-network expansion determines the amoeboid phenotype (22).

Evans and Yeung studied the small and large deformation of the cell passing through the micropipette to establish a theoretical model of the cell, the cell model is composed of a cortex wrapped with cytoplasm, where the cortex is regarded as anisotropic with static tension in the fluid layer and the cytoplasm is regarded as Newtonian fluid (15, 24). Theret studied the shear deformation of cells with micropipetting technology, mathematically modeling cells as incompressible elastic or viscoelastic solid homogeneous materials (25). Schmid-Schönbein studied the rheological properties of leukocytes with small deformation, and treated the cells as a homogeneous viscoelastic solid model (26). Skalak simulated the expansion and contraction of cells with thermodynamics and continuous mechanics, treating cells as a cytoplasmic gel structure containing actin, microtubules, and fiber filaments (27). Dong studied the nonlinear large deformation of cells by treating the cells as a layer of cortical layer solids with prestress and elastic tension containing Maxwell fluid (28), while Hochmuth only regarded the cells as a simple surface with constant cortical tension Newton’s droplet model to study the small deformation state of cells (29).

Experimental studies have found that when the cells protrude the bleb, the cytoplasm flows into the bleb, the cell volume does not change, and the cell membrane is separated from the actin skeleton at the bleb area. Many scholars conduct numerical simulation research on blebbing. Young and Mitran established a complex mathematical model. In the model, the cell structure is treated as a cytosol containing elastic Newtonian fluid and elastic fiber filaments (30). The cytoskeleton protrudes from the membrane to form a pseudopod. The disadvantage of this model is that the fiber filament will penetrate the cell membrane during the simulation process, and the effect on the membrane protrusion is unknown. Lim established a cell model composed of cell membrane, cortex and cytoplasm. The cells are set in a single channel to study the migration, the results showed that cell migration speed is affected by the size of the channel (31–32). Strychalski and Guy established a cell numerical model composed of elastic cell membranes and connecting bonds, porous elastic cortex and Newtonian fluid cytoplasm (33–34). The numerical experiments showed that the viscosity and drag of the cortex play a significant role on the blebbing dynamics. Woolley established the cell model composed of as an axisymmetric elastic shell surrounding incompressible fluid and weakened a small part of the shell to model the initiation of the bleb (35–36). With the mechanism of the membrane growth the model can produce the results that are of the correct quantitative size of the bleb comparing with the results of Charras’s (37).

This paper explores and studies the mechanical phenomena of amoeboid-type cell migration. A two-dimensional fluid-solid coupling cell model is established, and a numerical solution method for fluid-solid coupling is developed to simulate the behavior of cell bleb by releasing the connection between the membrane and the cortex of a cell. The effects of parameters such as the number of broken bonds, the viscosity coefficient of the cortex, and the cell membrane modulus on the size and shape of the cell bleb were investigated.

## 2. FLUID-SOLID COUPLING CELL MODEL

The structure of an actual animal cell is very complicated (as shown in Fig. 1A), it includes cell membrane, cortex, cytoplasm, nucleus and various organelles in macro-level. As reviews, there are different cell mechanics models to face various challenges. In our analysis, a two-dimensional cell model is recommended (Fig. 1B), each component is modeled as:

- The cell membrane is regarded as a linear elastic membrane with initial prestress;
- The cortex is a fiber network attached to the cell membrane which may sustain extension in both radial and tangential directions, therefore it can be simplified as a combination of a permeable elastic membrane (called cortex membrane) and elastic bonds between cell membrane and cortex membrane;
- The cytoplasm is regarded as a kind of imcompressible viscous fluid;
- The role of the nucleus is ignored and its mass is equivalent to the cytoplasm, due to the slow expansion of bleb.

**Figure 1.**
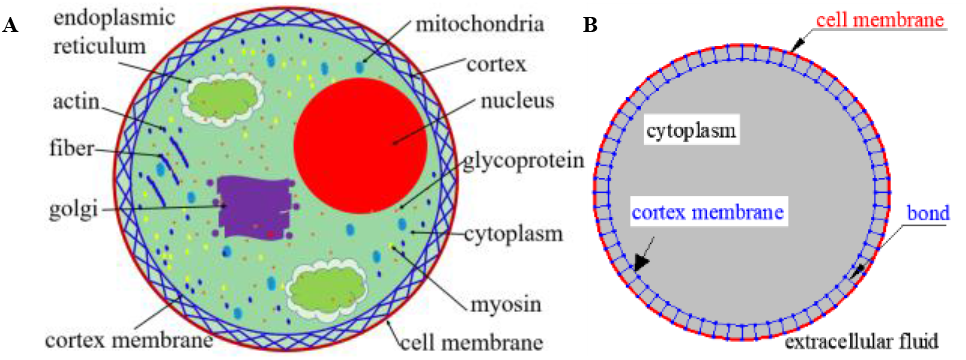
Cell structure. (A) The structure of a eukaryotic cell: contains cell membranes, cortex, actin, golgi, myosin, cytoplasm and so on; (B) Cell model: the two-dimensional fluid-solid coupled cell model established in this paper includes cell membrane, cortex, bonds, cytoplasm, extracellular fluid.

Obviously, the cell model in this paper is a fluid-solid coupling model.

## 3. GOVERNING EQUATION

### 3.1 Cytoplasm and extracellular environment fluid

Assuming that the cytoplasm and environmental liquid are incompressible Newtonian fluids, the governing equation of the flow field is the incompressible Navier-Stokes equation. The Euler coordinates are used to describe the motion of the fluid:

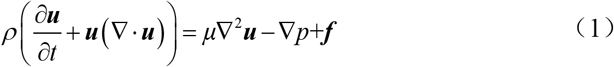

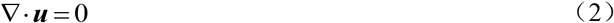

Where *σ* is the fluid density, ***u***(***x***,*t*) and *p* (***x***,*t*) are the fluid velocity and pressure respectively, ***x***(*x*, *y*) is the Euler grid point vector coordinate of the flow field, *μ* is the hydrodynamic viscosity coefficient, and ***f***(***x***,*t*) represents the volume force applied on the fluid.

### 3.2 Cell membrane and cortex membrane

The cell membrane and cortex membrane are elastic membranes with initial tension, therefore their governing equations are:

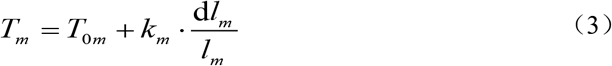

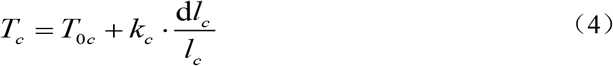

In which, *T_m_* and *T_c_* are the current tensions in cell membrane and cortex membrane respectively, *T*_0*m*_ and *T*_0*m*_ are the two initial tension in two membranes; *k_m_* and *k_c_* are the elastic coefficients of cell membrane and cortex membrane respectively; *l_m_* and *l_c_* are arc length of a section of cell and cortex membranes.

### 3.3 Bonds between cell membrane and cortex membrane

A bond between cell membrane and cortex membrane is regarded as a linear spring, its governing equation is,

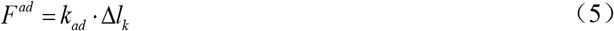

Where *F^ad^* represents the force between the cell membrane and the cortical membrane, Δ*l_k_* is the change displacement between the two membrane, *k_ad_* the stiffness coefficient of the linear spring.

### 3.4 Interaction between permeable cortex and cytoplasm fluid

Because of the cortex permeability, there is relative motion between cortex and cytoplasm. The interaction between the cortex and the cytoplasm fluid is regarded as a damping of a solid particle in fluid, which is proportional to their difference of velocity,

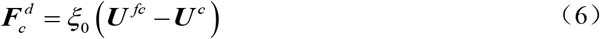

Where 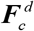 is the drag force, *ξ*_0_ is the damping coefficient of the fluid at the cortex, ***U***_*fc*_ and ***U***_*c*_ are the velocity of the fluid on the cortex and the velocity of the cortex

## 4. Discretization of Governing Equations

In order to solve the cell deformation under various conditions, the continuous solid-fluid coupling cell model is discretized (Fig. 2). In the discrete model, since the cytoplasm can permeate the cortex, the fluid-solid coupling between the cortex and the cytoplasm is described with a relative damping motion between the solid particles of the cortex and the fluid point in the same space. The solid points in the cell membrane and the fluid points in the same space are relatively stationary, since the cytoplasm and extracellular environment fluid can’t permeate the cell membrane.

**Figure 2.**
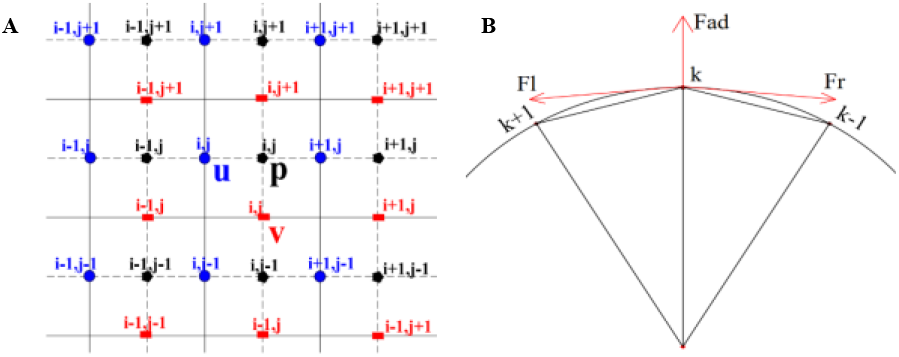
Discrete analysis. (A) Interlaced grid points: u, v, p represent fluid horizontal velocity, vertical velocity and pressure, respectively; (B) Stress analysis at the k-point on the membranes.

### 4.1 Discretization of fluid in Euler Space

The cytoplasm and the extracellular environment fluid are discretized using Euler space grids. As shown in Fig. 2A, a staggered grid will be used, where *u* represents the horizontal velocity of the fluid, *v* represents the vertical velocity of the fluid, and *p* represents the pressure of the fluid. Interpolate the grid points of horizontal velocity and grid points of vertical velocity to the grid point velocity where the pressure is located to avoid errors that may be caused by the second-order differential, which makes the calculated value more accurate. As shown in the Eq. 7 and Eq. 8:

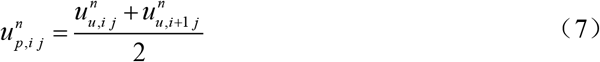

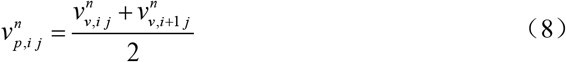

*n* represents the number of iteration steps, 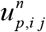, 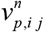 represents the horizontal velocity and vertical velocity of the fluid at the pressure grid point (*i*, *j*) when the iteration step is *n*; When the iteration step is *n*, 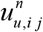 represents the horizontal fluid velocity at the horizontal velocity grid point (*i*, *j*); 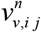 represents the vertical fluid velocity at the vertical velocity grid point (*i*, *j*).

Then substitute equation (2) into equation (1) to obtain the flow velocity of the iterative step *n*+1:

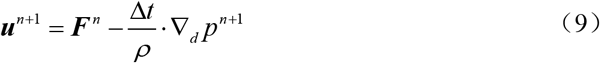

Where ***F***^*n*^ is the result of iteration step *n*

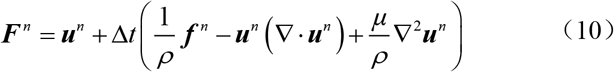

Taking the divergence of Eq. 7, the pressure field of the fluid in *n*+1 iteration steps can be obtained

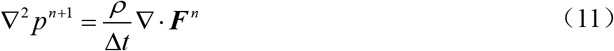

The equations (10) and (11) are analyzed by the finite difference method to obtain the pressure field *p* in the fluid domain, and then the pressure field *p* is brought into the equation (9) to obtain the velocity field ***u*** in the fluid domain.

### 4.2 Discretization of membranes in Lagrange Space

The cell membrane and the cortex are divided into *N* discrete points in Lagrangian coordinates system. Force analysis is performed on solid discrete nodes (Fig. 2B). The discrete node *k* is subjected to the stretching effect of the discrete point on the left *k*+1 and the *ad* discrete point on the right *k*−1 to this discrete node, thereby the tension on the discrete node *k* is defined as follows:

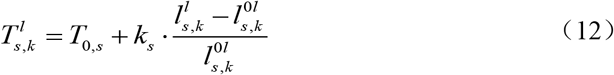

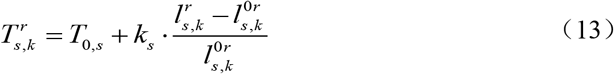

where, the subscript *s* = *m*, *c*, *m* is the cell membrane and *c* is the cortex 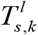 and 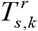 respectively represent the magnitude of the tension on the discrete point *k* by the discrete point on its left and the discrete point on its right; *T*_0,*s*_ represents the initial solid tension; *k_s_* means solid stiffness coefficient; 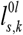 and 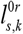 respectively indicate the distance between the discrete point *k* and its left discrete point, and the distance between the discrete point and its right discrete point at the initial iteration step *t*_0_; 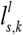, 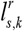 respectively indicate the distance between the discrete point *k* and its left discrete point, and the distance between the discrete point and its right discrete point at the time of the iterative calculation.

The solid discrete point *k* is subjected to concentrated loads applied by the left and right discrete point, respectively:

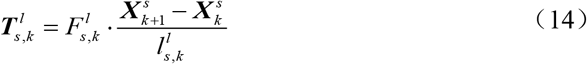

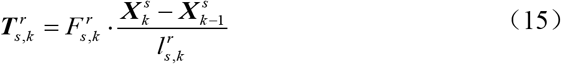

Where 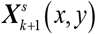, 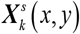 and 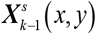 are the Lagrangian vector coordinates of the solid discrete points *k*+1, *k* and *k*−1, respectively. 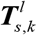, 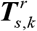 are the discrete point of the solid subject to the tension of the left and right discrete point, respectively.

The bond connect the node *k* of the cell membrane and the node *k* of the cortex membrane is regarded as a linear spring. The concentrated loads of the bonds applied on the node *k* of the cell membrane and the cortex membrane have the same magnitude and opposite direction:

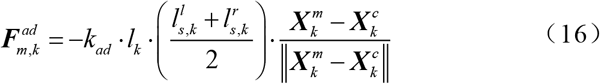

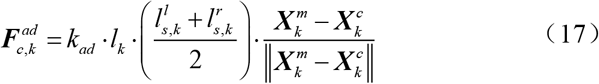

where, *k_ad_* is the elastic coefficient of the connecting bond; 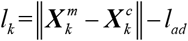 is the deformation of bond between the cell membrane and the cortex at discrete point *k*, *l_ad_* is the initial length of this bond; 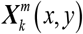, 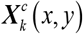 are the Lagrange discrete points of the cell membrane and the cortex vector coordinates. 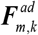, 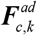 are the equivalent concentrated load on the node of the cell membrane and the cortex membrane respectively apply by the bond between them.

The resultant force at the discrete point *k* of the cell membrane is:

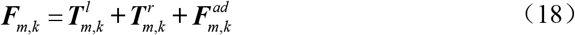

The resultant force at discrete point *k* of the cortex membrane is:

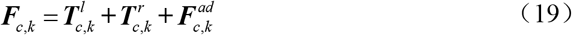

### 4.3 Fluid-solid coupling mutual conversion

The immersion boundary method is a mathematical modeling method as well as a numerical discrete method (38–40). The effects of solid on the fluid at the boundary are transformed into discrete body forces in the flow field, which is characterized by using Euler and Lagrangian variables to mark the position of the flow field calculation point and the immersion boundary respectively. In this paper, the Euler variable of the fluid is discretized into a square grid in the cartesian coordinate system, and the Lagrange variable of the solid is described in the polar coordinate system.

#### (a) Volume forces applied in fluid

The unit volume force 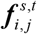 on the Euler fluid grid point (*i*, *j*) at the current moment *t* is calculated according to the current position and force of the Lagrangian discrete point *k* of the immersed boundary. The distribution equation of 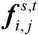 is:

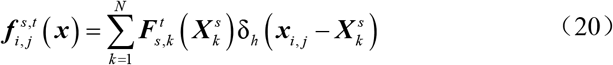

#### (b) Convert fluid velocity to solid velocity

The velocity 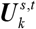 at the discrete point of the immersion boundary can be obtained by interpolation of the velocity of the flow field near the current moment, and its distribution equation is:

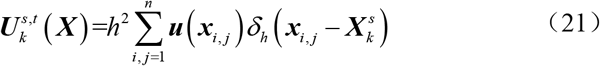

Where *h* is the grid size of fluid, and *δ*_h_ represents the regularized *δ* function, which converts variables between solids and fluids. The general two-dimensional formulation can be written as follows:

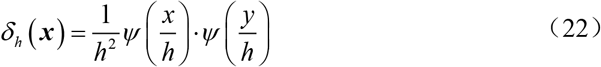

Where *ψ* can have multiple forms, the following form is selected (39):

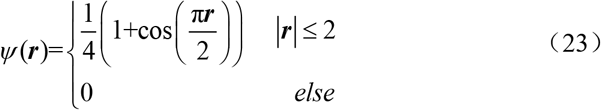

The slip-free boundary condition is applied to the cell membrane on the immersion boundary, the expression is 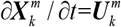. Because the cortex is permeable, it affects the speed of the cortex when cytoplasm flows. According to the equilibrium conditions, the resistance at each discrete point on the cortex is 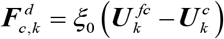, *ξ*_0_ is the damping coefficient of the fluid at the cortex, 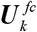 and 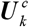 are the velocity of the fluid on the cortex and the velocity of the cortex at the discrete point *k*.

## 5. NUMERICAL SOLUTION PROCESS

### 5.1 Initial condition and boundary condition

The cell is set in a stationary flow field with a size of *L_x_* ∗ *L_y_*. The domain is divided into *n_x_* ∗ *n_y_* fluid grid. The cell membrane and the cortex are divided by *N* discrete mark points, and connected at the mark points by the bonds. The pressure boundary condition of the extracellular flow field is *p_out_* = 0, slip free boundary conditions were applied to the cell membrane.

### 5.2 Material parameters

The neutrophil bleb is studied with the developed numerical method. The material parameters used are shown in Table 1, and the grid parameters are shown in Table 2.

**Table 1.** Material parameters

| Description | Symbol | Value | Quote |
| --- | --- | --- | --- |
| membrane radius | $r_M$ | 10 $\mu\text{m}$ | (6) |
| cortex radius | $r_C$ | 9.0625 $\mu\text{m}$ | (32) |
| initial membrane tension | $T_{0m}$ | 40 pN/$\mu\text{m}$ | (6) |
| initial cortex tension | $T_{0c}$ | 250 pN/$\mu\text{m}$ | (6) |
| membrane stiffness coefficient | $k_{sm}$ | 4 pN/$\mu\text{m}$ | (44) |
| cortex stiffness coefficient | $k_{sc}$ | 100 pN/$\mu\text{m}$ | (6, 37) |
| membrane-cortex spring constant | $k_{AD}$ | 29.5 pN/$\mu\text{m}^3$ | |
| viscosity of fluid | $\mu$ | 0.003~2 pa·s | (41-42) |
| cortical damping coefficient | $\xi_0$ | 10~10<sup>4</sup> pN/$\mu\text{m}^3$ | (34) |
| cortex permeability coefficient | $\xi$ | 10~300 pN/$\mu\text{m}^3$ | (34) |

**Table 2.** Calculation parameters

| Description | Symbol | Value |
| --- | --- | --- |
| solid grid discrete points | $N$ | 256 unit |
| fluid grid discrete points | $n_x, n_y$ | 64~128 unit |
| fluid domain range | $L_x * L_y$ | 30*30~60*30 $\mu\text{m}^2$ |
| time step | $dt$ | 0.001 s |

### 5.3 Flowchart of numerical solution

The solution process is shown in Fig. 3 below:

**Figure 3.**
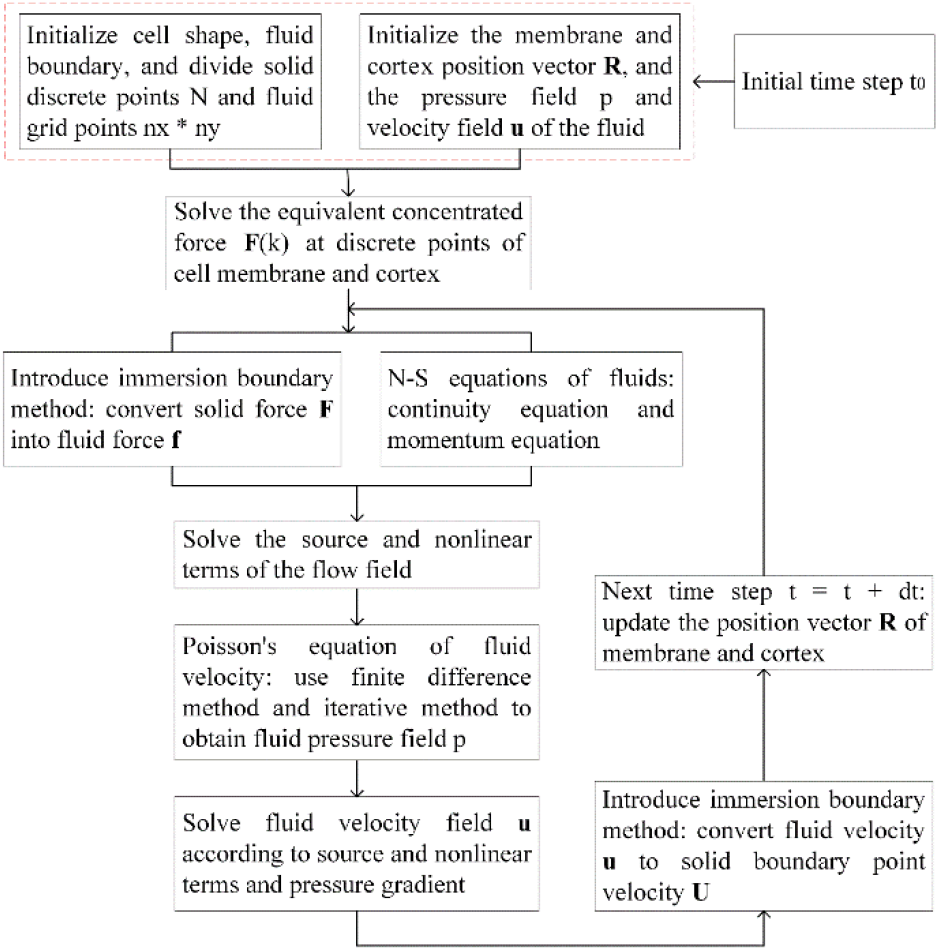
Flowchart of numerical solution

## 6. RESULTS AND ANALYSIS

The first numerical analysis of a period of 30s is carried out to study the stability of the cell model when there is no bleb generation. The second numerical analysis is carried out to simulate the cell bleb growth process of a period 30s; we first let the model stand for 1s, then break the bonds to calculate 29s to observe the growth process of the bleb.

### 6.1 Stability analysis of the model

Various the characteristic parameters of cell are considered to study the reliability of the numerical model. By changing the viscosity of the cell fluid, the damping coefficient of the cortex, and the value of the cell membrane-cortex elastic modulus, the effects of these changes on fluctuation in the cell flow field and cell shape were explored, and the rationality and stability of the cell model were analyzed.

#### 6.1.1 No-bleb

##### (a) Viscosity of the cell fluid

According to experiments (41–42), the cytosol of different cells have different dynamic viscosities, ranging from 0.01 Pa·s to 100 Pa·s. Numerical simulations were performed for different viscosities of the cytosol, and the maximum velocity value of the flow field and the maximum velocity increments of the contiguous time steps, as well as the changes in the shape of the cell membrane and cortex over time were analyzed. As shown in Fig. 4A, as the time increases, the greater the viscosity leads to the smaller the maximum velocity value of the flow field; the maximum velocity value of the flow field decreases with time and tends to be stable (the maximum velocity increments of the contiguous time steps gradually approaches zero), which indicates that the larger the viscosity, the earlier it reaches the stable state. For different cell fluid viscosities, the pressure of fluid inside and outside the cell remains basically unchanged, and the maximum pressure in the cell is about 29 Pa. This pressure can achieve an equilibrium state with the solid internal forces. The results showed that the present numerical model can simulate the cells with different cytosol viscosities.

**Figure 4.**
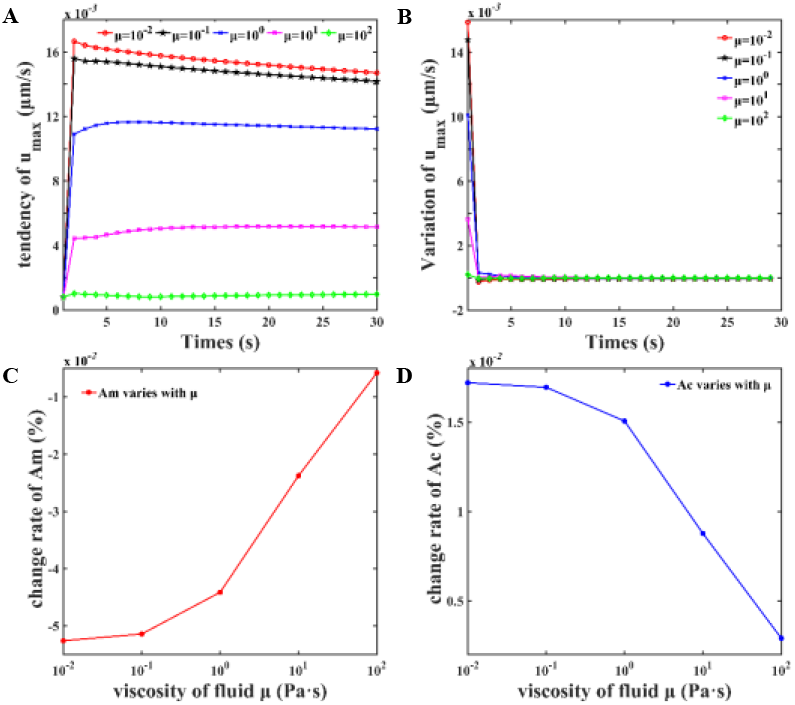
Change the viscosity of the fluid. (A) The maximum value of the flow field velocity; (B) The time step of the maximum velocity change with time; (C) Cell membrane area change rate with the viscosity; (D) Cortex area change rate with the viscosity.

The change rate of the cell membrane area is defined as

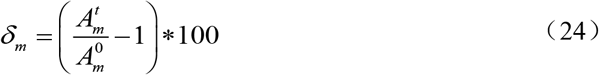

Where 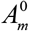 is the area at the initial moment of the cell membrane, and 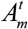 is the area at the final moment of the cell membrane. As shown in Fig. 4B, its absolute value ranges from 0% to 0.06%. The change rate of the cortical area is defined as:

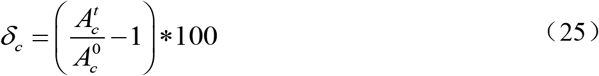

Where 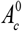 is the area of the cortex at the initial moment and 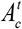 is the area of the cortex at the final moment. As shown in Fig. 4C, it ranges from 0% to 0.02%. It can be obtained that the shape of the cell remains basically unchanged, and it means that the cell is in a stable state. The shape change of the cells is almost negligible, which means the numerical model can simulate the balanced and stable state of cell with various dynamic viscosities.

##### (b) Damping coefficient of the cortex

The cortex is polymerized by actin, and actin is reassembled according to environmental changes, so the damping coefficient of the cortex varies widely (43). The value of the cortex damping coefficient represents the ability of the cytoplasm to penetrate the cortex, which has significant effect on the flow of intracellular cytoplasm, and then changes the shape of cell during the bleb formation. In this paper, damping coefficients of different orders of magnitude are selected between 10 pN·s/μm^3^ ~10^4^ pN·s/μm^3^. As shown in Fig. 5A, for different cortical damping coefficients, the maximum fluid velocity values of the cell remain basically the same, which are about 0.0108 μm/s, and the increment of the maximum velocity values tends to zero with time, which means that the influence of the damping coefficient of the cortex on the flow field can be ignored for the stable cell. Meanwhile, the maximum values of the cell pressure of fluid remain basically unchanged at 29 Pa·s consistent with the equilibrium state. As shown in Fig. 5B and Fig. 5C, the absolute value of the change rate of the cell membrane area ranges from 0.043% to 0.046%, and the absolute value of the change rate of the cortex area ranges from 0% to 0.02%, which means there is almost no change in the cell shape.

**Figure 5.**
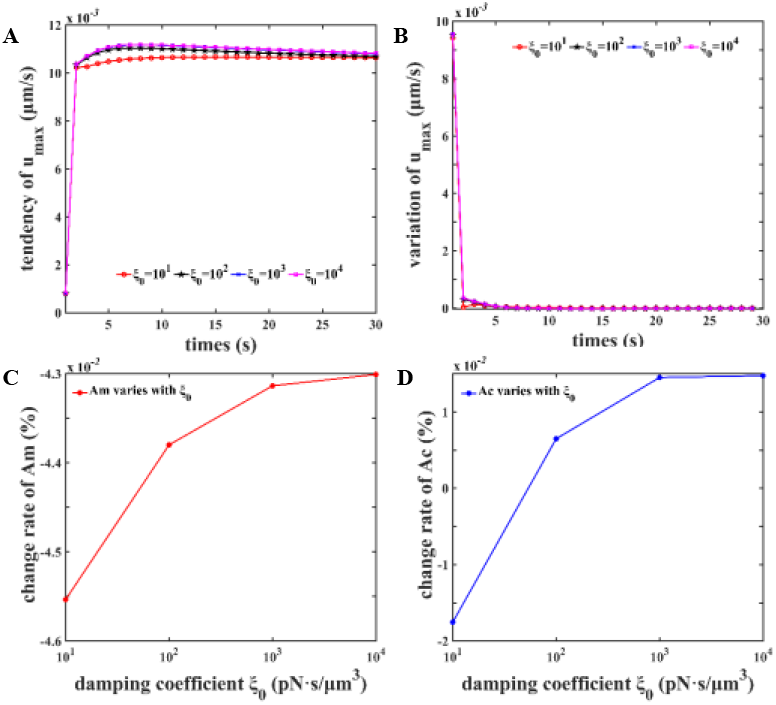
Change the damping coefficient of the cortex. (A) The maximum value of the flow field velocity; (B) The time step of the maximum velocity change with time; (C) Cell membrane area change rate with viscosity; (D) Cortex area change rate with viscosity.

The results showed that the present numerical model can simulate the cells of stable state with a wide range of cell cortex damping coefficients 10 pN·s μm^3^ ~10^4^ pN·s μm^3^. It is considered that the present model is stable for the damping coefficient in this range.

##### (c) Value of the membrane and cortex elastic modulus

The elastic modulus of the cell membrane and the cortex (1):

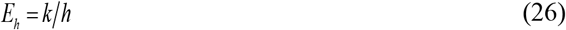

Where *k* is the stiffness coefficient (unit: pN/μm) and *h* is the solid thickness (unit: μm). Experiment measured that the thickness of the cell membrane was about 0.1 μm, and the thickness of the cortex was also about 0.1 μm. According to the literature, the different stiffness coefficient values of the cell membrane and the cortex are selected for numerical experiment. In this chapter, we take the cell fluid viscosity *μ* = 1.2 Pa·s and the damping coefficient *ξ*_0_ =1.1e4 pN·s/μm^3^. The numerical simulations are carried out with various stiffness coefficients of the cell membrane and the cortex. The experimentally measured cell elastic modulus of softer cells (blood cells) is about 1 Pa, and the cell elastic modulus of stiffer cells (endothelial cells) is about 1000 Pa (44). The elastic modulus of the cell is mainly determined by the mechanical properties and structure of the cell membrane and cortex, but the elastic modulus of the cortex of the medium-strength cells measured by Tinevez is about 240 Pa, and the cortex has a strong ability to change (6), so we set the elastic stiffness (*k_m_*, *k_c_*) of the cell membrane and the cortex is changed from (2,10^0^) to (10,10^3^). The results are shown in Fig. 6:

**Figure 6.**
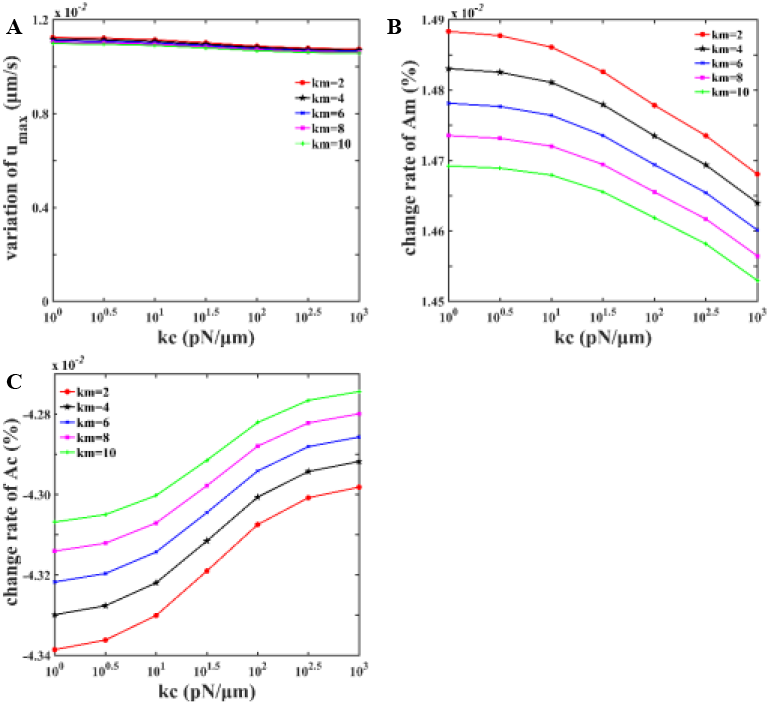
Change the elastic modulus of cell membrane and cortex. (A) At t = 30s, maximum change of flow field velocity under different elastic modulus of cell membrane and cortex; (B, C) Cell membrane and cortex area change rates change with the elastic modulus of the membrane and

Fig. 6A shows that the maximum velocity of the corresponding flow field for different elastic modules at 30s is basically the same; Fig. 6B shows that for different elastic modulus combination, the errors of the cell membrane and cortical area change rates are 0.0004% and 0.0007%, which means the shape of the cells remains unchanged. The influence of the elastic modulus of the cell membrane and the cortex on the cell shape is basically negligible, which proves the correctness and stability of the cell model within the range of the elastic modulus of the cell membrane and the cortex.

#### 6.1.2 Bleb simulation

According to experimental observations (45), changing the cortex-related mechanical properties can lead to the generation of bleb in cells. The internal and external fluid pressure fields and the solid structure keep an equilibrium state for stable cell. Intracellular pressure is the main reason to drive the cell membrane to expand outward and form the bleb (37, 46). Myosin contraction can cause local breakdown of the actin cortex, pressure drives the cytoplasm to flow out of the cortex, and then fill the cell gap between the membrane and the cortex, simultaneously expand the cell membrane to form the bleb (11). In order to study the effect of spontaneous cortex contraction on cell movement, many scholars have performed a variety of experiments to explain the specific reasons for multiple migration modes (14, 19). There are two main mechanisms to generate bleb: local decrease in membrane-to-cortex attachment, or local rupture of the cortex itself (13, 47). Because the cortex rupture is too small to image, these two mechanisms are combined in the generation of bleb (13).

In order to verify that this model can produce bleb same to the experimental results, as shown in Fig. 7. According to the experiment of Tinevez et al. (6), we changed the initial tension of the cortex and broke the bonds between the cell membrane and the cortex membrane consistent with the experimental treatment (6). In order to simulate the experimental drug treatment results, different connection bonds were broken for different initial tensions to get different arc lengths. The specific values are shown in the table 3.

**Figure 7.**
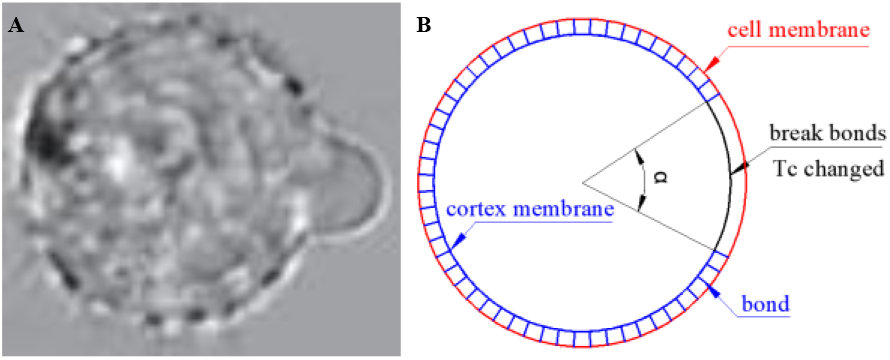
Bleb formation. (A) Bleb formation after changing the initial tension of the cortex (6); (B) Schematic diagram of the cell structure that breaks the bonds and changes the initial tension of the cortex.

**Table 3.** the specific values

| initial tension of<br>the cortex (6)<br>pN/$\mu m$ | the arc length of<br>the bonds for<br>the hole $\mu m$ | range point of<br>broken bonds |
| --- | --- | --- |
| 100 | 2.364 | 6~252 |
| 200 | 3.256 | 8~250 |
| 300 | 4.166 | 9~249 |
| 400 | 5.134 | 11~247 |
| 600 | 5.968 | 13~245 |

Because the change range of the viscosity of the cell fluid is wide, we set the specific values as: 0.0012, 0.012, 0.12, 1.2 (unit: Pa·s) to perform numerical calculations of 30s is performed on this. As shown in Fig. 8, when the initial tension of the cortex is 500 pN/μm, the morphology of cells and bleb at different viscosities, the phenomenon of cells generating bleb is close to the experimental results (6).

**Figure 8.**
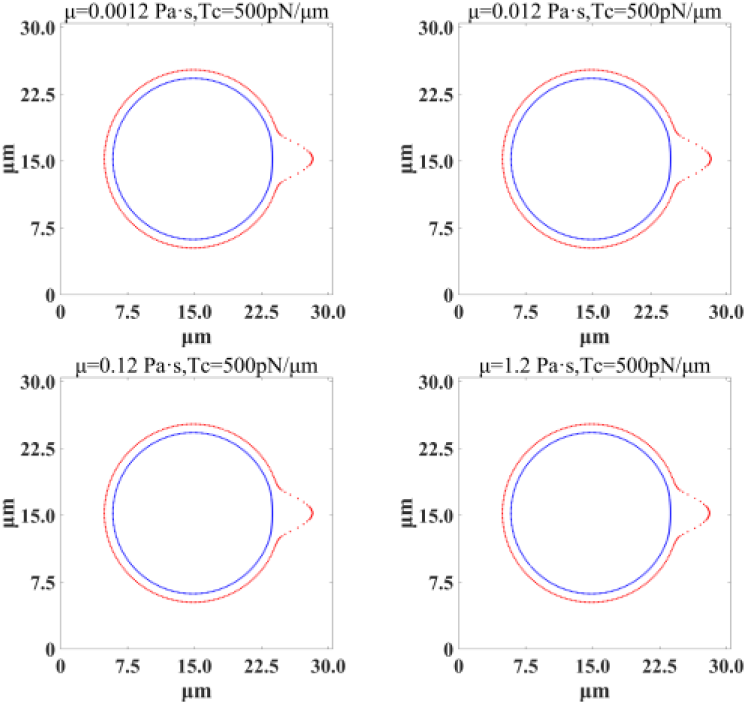
Bleb formation. When the initial tension of the cortex is 500 pN/μm, the growth morphology of blebs under different cell fluid viscosity at t=30s.

Comparing the growth of the bleb protrusion with the experimental results, as shown in Fig. 9, the viscosity of the cell fluid has little effect on the morphology of bleb. Due to the symmetry of the cell model, the horizontal displacement of the first point on the cell membrane is defined as the length of the bleb protrusion, and the change in the arc length of the broken bonds is defined as the width of the bleb. As shown in Fig. 9A, the length of the bleb extending from 0.5 μm to 2.8 μm, the experimental result is between 1.8 μm to 3.5 μm, the width of the bleb varies from 1.2 μm to 2.6 μm, as shown in Fig. 9B, the experimental numerical result is 0.8 μm ~3 μm (6).

**Figure 9.**
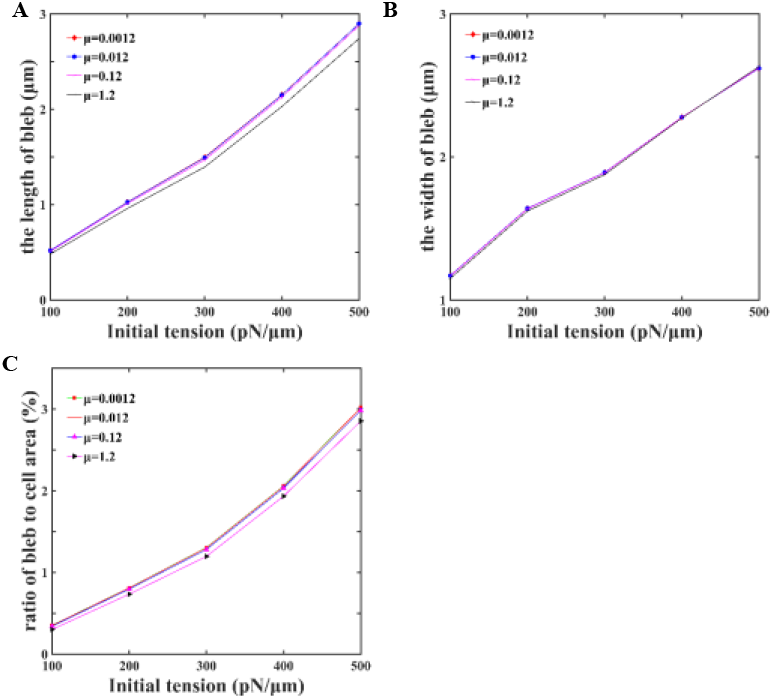
Bleb analysis that changes the initial tension of the cortex at different cell fluid viscosities. (A) The length of bleb; (B) The width of bleb; (C) The area ratio of bleb.

In summary, when the initial tension of the cortex is less than 200 pN/μm, the cells can only generate very small bleb, and the blebs of the cells are not obvious. The ratio of blebs to the cell area changes with the different initial tension of cortex as shown in Fig. 9C. Therefore, the results shown that this model can predict the experimental results of blebs growth, and the cell model is also reasonable and reliable.

### 6.2 Discussion about the bleb

According to experimental results, the angles *α* of the local fracture area of the membrane-cortex are selected in this paper as: 30 °, 45 °, 60 °, and 90 °, respectively (Fig. 10). Then the numbers *N*_0_ of membrane-cortical bonds that need to be broken for the corresponding arc length are 22, 32, 42, 64 (unit: unit). On the other hand, it is found from Fig. 5C that the cortex has contractility in the range of 10 pN·s/μm^3^ ~10^2^ pN·s/μm^3^, but the specific damping coefficient that affects the permeability of cells is unknown. Therefore, the values of the local damping coefficient *ξ* of the cortex in the key broken area are: 10, 20, 35, 50, 75, 100, 150, 200, 300 (unit: pN·s/μm^3^). The bleb growth process simulation was performed on different combinations of cell membrane-cortex bonds area angle *α* and different local damping coefficient *ξ*.

**Figure 10.**
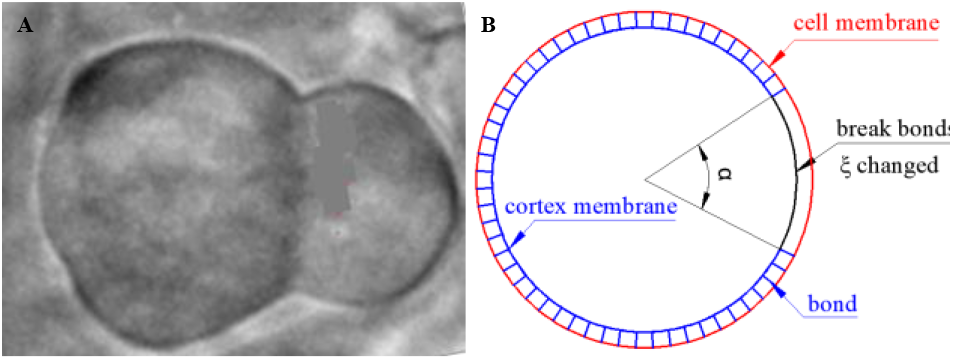
Bleb simulation of large cell deformation. (A) Bleb phenomenon produced in the experiment (11); (B) Schematic diagram of cell structure with changing the fracture angle and local damping coefficient of cortex.

#### 6.2.2 Analysis of flow field

As shown in Fig. 11 and Fig. 12, the state of the flow field within 60 s is analyzed when the fracture angles are 30 °, 45 °, 60 ° and 90 ° with different local damping coefficients of cortex.

**Figure 11.**
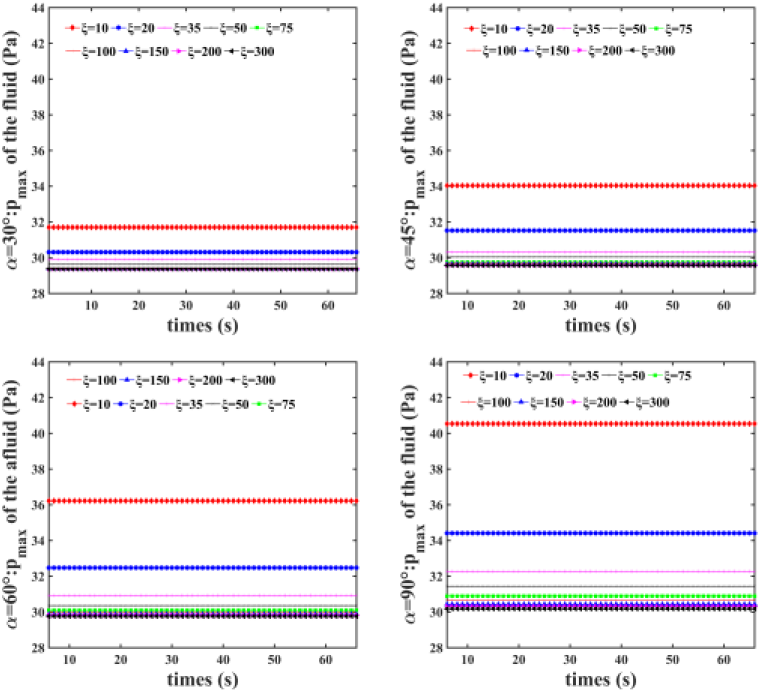
When the fracture angles are 30 °, 45 °, 60 °, and 90 °, the maximum value of the flow field pressure changes within 60 s under different local damping coefficients

**Figure 12.**
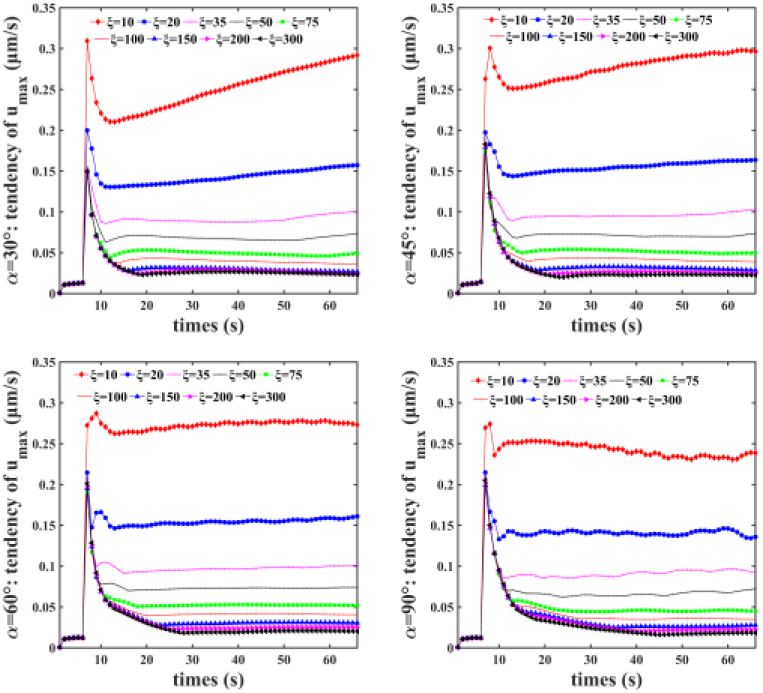
When the fracture angles are 30 °, 45 °, 60 °, and 90 °, the maximum change of the flow field velocity within 60s under different local damping coefficients

The simulation results show that for all the fracture angle considered, the maximum pressure value in the flow field remains stable and is close to about 30 Pa, which is within the range of the pressure measured in the experiment. But as the fracture angle increases, the pressure in the flow field will increase. On the other hand, as the local damping coefficient increases, the maximum fluid pressure value gradually decreases. When the damping coefficient is less than or equal to 20 pN·s/μm^3^, it will have a greater impact on the pressure; when the local damping coefficient is greater than 35 pN·s/μm^3^, the pressure will remain basically constant, about 30 Pa. For the velocity of the flow field, the larger the fracture angle, the faster the maximum velocity of the flow field ***u***_*max*_ decreases. When the fracture angle is greater than or equal to 60 °, the maximum velocity ***u***_*max*_ of the flow field stabilizes faster. As the local damping coefficient increases, the faster the maximum velocity of the flow field ***u***_*max*_ tends to zero, the faster the flow field can reach a relatively stable state.

As shown in Fig. 13A, the pressure value increases with the increase of the fracture angle *α*, but when the damping coefficient *ξ* is greater than or equal to 75 pN·s/μm^3^, the pressure value is nearly constant regardless of the increase of the fracture angle *α*. As shown in Fig. 13B, the magnitude of the flow velocity depends on the damping coefficient *ξ*. The smaller the damping coefficient leads to the greater flow velocity. But for the damping coefficient *ξ* lager than a certain value, the velocity of the flow field is basically the same.

**Figure 13.**
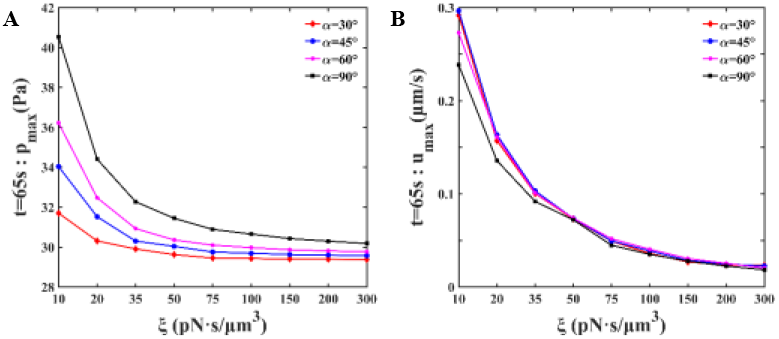
Changes in cell flow field. (A) The maximum pressure value of the flow field varies with the fracture angle and local damping of the cortex; (B) The maximum velocity of the flow field varies with the fracture angle and local damping of the cortex.

#### 6.2.2 Analysis of bleb morphology

##### (a) The pressure of the bleb

For the parameters combination *α* =90 °and *ξ* = 20 pN·s/μm^3^, the fluid pressure and the shape of the cell at different times are shown in Fig. 14. The pressure field in the cell can achieve an equilibrium state with the solid structure, and the change in cell shape is basically the same as that in the experimental phenomenon of the amoeboid type of migration cell during the cell pseudopod growth process (11). The results show that the angle *α* and the damping coefficient *ξ* of the cortex at the broken bonds can determine the contraction of the cortex, thereby affect the speed of the cytoplasm flows from the inner region of the cortex to space between the cortex and the cell membrane, and then determine the speed and size of bleb growth. When t = 50s, the pressure field inside and outside the cell can reach a stable state, a period of 60s after the broken bonds is carried out to simulate the bleb growth.

**Figure 14.**
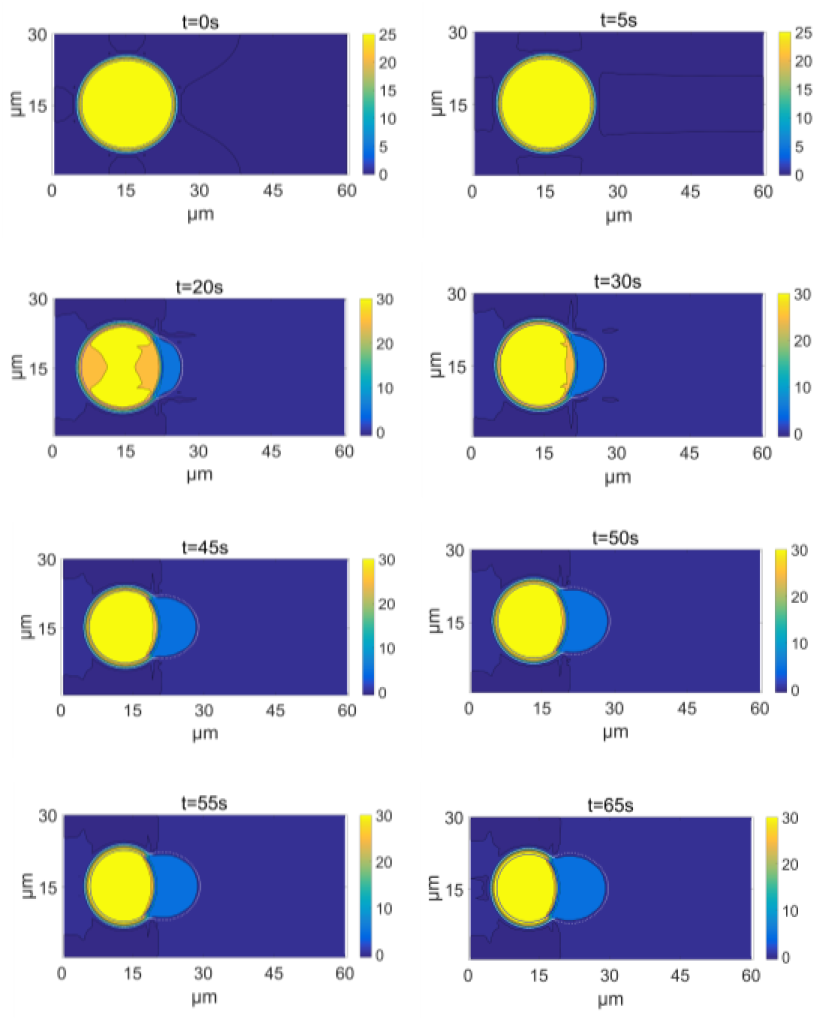
When the fracture angle is 90°, and the local damping coefficient is 20 pN·s/μm^3^ and cell shape at different times.

##### (b) Velocity of the bleb

As shown in Fig.15, it can be seen from the figure that when the fracture angle is smaller, the time required for bleb growth to stabilize is shorter; and when the fracture angle is greater than 60 °, the bleb growth rate results tend to be consistent. For different fracture angles, the bleb velocity changes greatly when the damping coefficient is less than or equal to 20 pN·s/μm^3^, reflecting the faster bleb growth and larger deformation. When the local damping coefficient is greater than 20 pN·s/μm^3^, the shorter the time required for the bleb velocity to stabilize.

**Figure 15.**
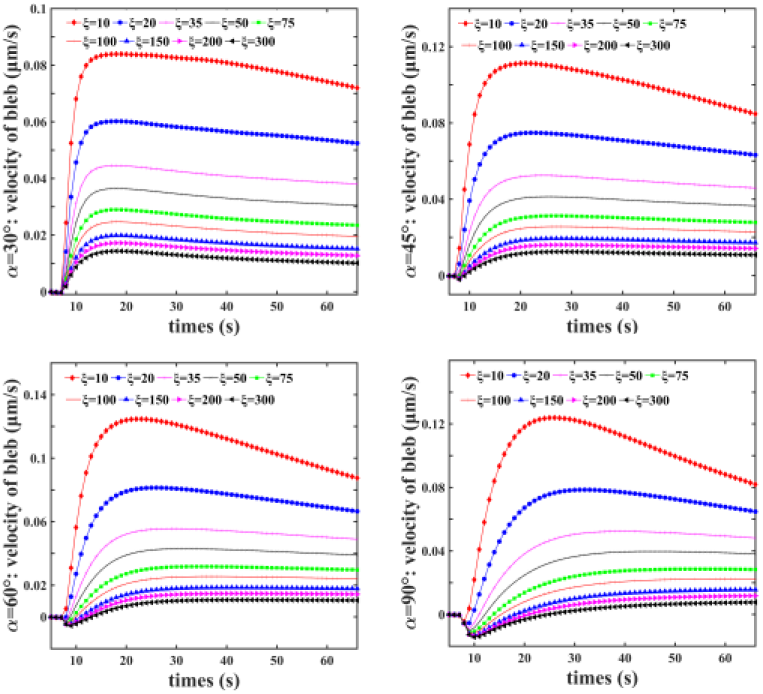
When the fracture angles are 30 °, 45 °, 60 ° and 90 °, the horizontal velocity of the bleb changes within 60s under different damping coefficients

According to the actual situation, when the bonds is suddenly broken, the flow field near the local cell membrane and cortex corresponding to it will produce a certain oscillation, then relatively speaking, the bleb velocity will have a certain oscillation within a few seconds after the bonds is broken. Therefore, when the fracture angle is 90 °, the bleb growth will be more in line with the real situation.

The change of the bleb length *l_b_* is shown in Fig. 16A. The bleb extension length is not affected by the fracture angle, but it will decrease with the increase of the local damping coefficient, and the extension length ranges from 0.5 μm to 6 μm (48). Therefore, it can be considered that the result is correct under the local damping coefficient within a certain range. Then, the relationship between the bleb extension speed and the local damping coefficient can be calculated. As shown in Fig. 16B, the extension speed decreases *v_b_* with the local damping coefficient between 0.01 μm/s and 0.9 μm/s. At the same time, it can be seen that the size of the fracture angle has little effect on the bleb extension speed. When the fracture angle is greater than 30 °, the bleb extension speed remains basically the same.

**Figure 16.**
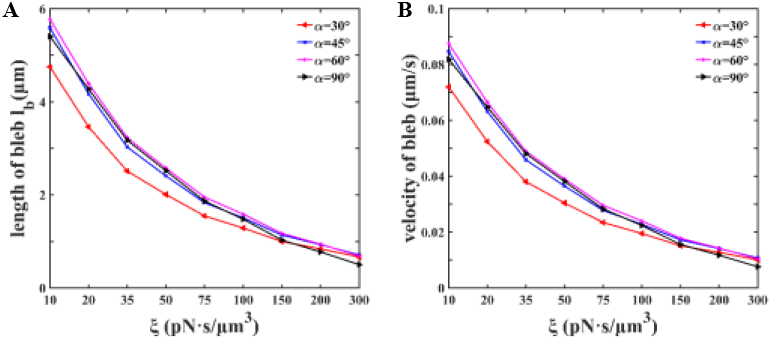
Bleb growth. (A) At different fracture angles, the length of bleb extension varies with the local damping coefficient of the cortex; (B) the bleb velocity varies with the local damping coefficient of the cortex.

##### (c) Area analysis of cell membrane and cortex

As shown in Fig. 17, different numbers of membrane-cortex elastic bonds are broken in the local area of the cell, and the damping coefficient of the cortex in the broken area is changed to simulate the process of the cell extending the bleb.

**Figure 17.**
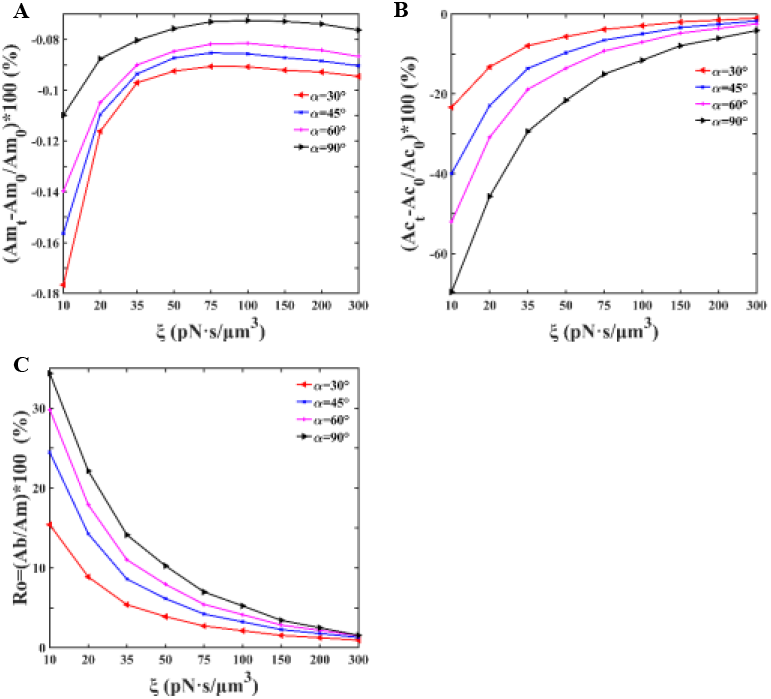
Changes in cell structure. (A) Cell membrane area change rate changes with the fracture angle and the local damping coefficients of cortex; (B) Cortex area change rate with the fracture angle and the local damping coefficients of cortex. (C) The change of the bleb area to the total cell area changes with the fracture angle and the local damping coefficient of the cortex.

As shown in Fig. 17A, the cell membrane area change rate is less than 0.2%, which means the cell membrane area is almost unchanged, the results are consistent with the facts. As shown in Fig. 17B, the cortex exhibits a larger contraction phenomenon, which is consistent with the experimental phenomenon (11), the larger the fracture angle *α* and the smaller damping coefficient *ξ*, the greater shrinkage of the cortex area, which is reasonable since the smaller *ξ* make the cytoplasm permeate through the cortex easily. When *α* = 90 ° and *ξ* = 10 pN·s/μm^3^, the cortex can reduce to 30% of the initial state.

According to the numerical simulation in the governing equations, the fluid pressure drives the flow in cell, and then protrude the bleb. The area ration of the bleb area to the total area of the cell is defined as:

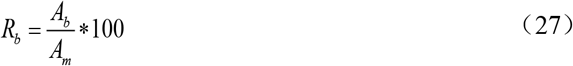

The bleb shape change of the numerical results and the bleb growth process in experiments were compared. As shown in Fig. 17C, the bleb area ratio increases with the increase of the angle of the ruptured membrane-cortex connection area, and decreases with the increase of the local damping coefficient. The area ratio *R_b_* ranges from 2% to 35%, Blaser measured the bleb volume between 0% and 8% (11); and the measured value of Tinevez is between 0% and 3% (6), which indicated that the present numerical method can simulate the bleb production of most cells with different shapes. Therefore, the present numerical model can simulate the generation of bleb with proper combination of fracture cell membrane-cortex connection area angle *α* and a local damping coefficient *ξ*.

In conclusion, the angle *α* of the local area of the rupture membrane-cortex elastic connection and the local damping coefficient *ξ* of the cortex at the broken bonds are two basic variables that affect the growth process of bleb. Define the size of the cortical contraction intensity *S_c_* as the reciprocal of the area change rate, that is:

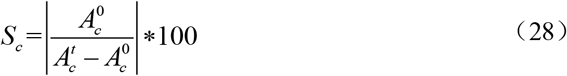

We can know from the previous chapter, the bleb extension speed *v_b_* is positively related to changes in the flow field pressure *p* and the fluid velocity ***u***. And then, the larger pressure *p* and the flow field velocity ***u***, the greater the bleb extension speed *v_b_*. According to the analysis, the local area *α* of the rupture membrane-cortical elastic connection and the damping coefficient *ξ* of the cortex at the broken bonds determine the contraction intensity *S_c_* of the cortex, while the contraction intensity *S_c_* of the cortex affects the flow field velocity ***u***, and then changes the flow field pressure *p*. The growth ability of the bleb is reflected by the bleb growth speed *v_b_*. From the analysis, it can be seen that the main factor affecting the bleb growth rate *v_b_* is the contraction intensity *S_c_* of the cortex, which is positively correlated and is expressed by Eq. 29

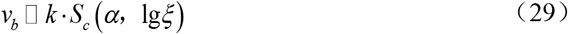

### 6.3 Parameters affecting bleb

Local decrease in membrane-to-cortex attachment results in a decrease of fluid pressure due to the balance of the cell membrane, which leads the cytoplasm inside the cortex flowing to the space between the cortex and cell membrane through the rupture area of the cortex. Because the cortex is formed by the polymerization of actin, the inter-molecular gap results in a certain permeability of the cortex. When a fibrous obstacle is encountered, the local cortex morphological change, such as local rupture and local permeability reduce of protein molecules between the membrane and the cortex, are induced by biochemical reactions, which will form bleb to avoid obstacle. It can be seen that the growth of the bleb is strongly related to the size of the local separation area and the permeability of the cortex. This article mainly studies the bleb growth process with a combination of the two mechanisms, weakening the structural role between the cell membrane and the cortex of the corresponding region by breaking local protein bonds, which increases the local permeability of the cortex to equal the local cortex rupture. A period of 60s is simulated to study the bleb growth.

The changes of cell shape, tension of the leading and trailing edges of the cell are further analyzed and studied. The influence of the fracture angle and the local damping coefficient on the cells generating bleb is discussed.

#### 6.3.1 Fracture angle

When the local damping coefficient is 20 pN·s/μm^3^, the 1^st^ point located at the leading edge of the cell and the 129^th^ discrete point at the trailing edge of the cell are selected to observe the effect of the fracture angle on the changes in cell membrane and cortex membrane tension.

As shown in Fig. 18, the smaller the fracture angle leads to larger tension increase of the cell membrane at the leading edge. The reason is that the initial cell membrane and cortex are in a state of tension. When the broken bonds are smaller, the fracture angle are smaller, and then the larger change that can be shared by the cell membrane or cortex. This explains that due to the partial loss of the bond’s constraint on the cell membrane, the local balance is destroyed, and the internal pressure expands the cell membrane and generates bleb. At the same time, during the bond break, the posterior edge of the cell membrane is far away from the destruction area, and it still maintains a balanced state, and the tension almost does not change. The tension of the anterior edge of the cortex membrane has a larger change, showing obvious contraction behavior; the tension of the posterior edge of the cortical membrane has a smaller change and has a smaller amplitude of contraction behavior. The swelling phenomenon of the cell membrane and the contraction behavior of the cortex are basically consistent with the experimental observation (12).

**Figure 18.**
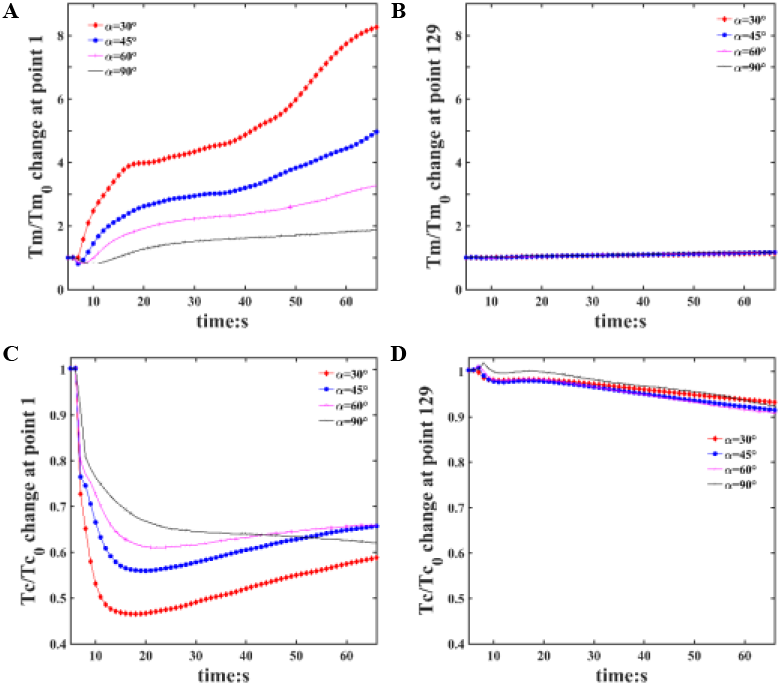
Changes in cell membrane and cortex tension. (A, B) When the fracture angles are 30 °, 45 °, 60 ° and 90 °, changes in tension of the cell membrane at 1^st^ point on the leading edge and 129^th^ point on the trailing edge; (C, D) When the fracture angles are 30 °, 45 °, 60 ° and 90 °, changes in tension of the cortex membrane at 1^st^ point on the leading edge and 129^th^ point on the trailing edge.

#### 6.3.2 Damping coefficient of the cortex

In this model, the membrane-cortex elastic bonds of a local area are broken and the cortex permeability of the local areas of is changed to simulate the change caused by actin polymerization in the cortex. These changes promote the flow of cytoplasm in the cell body, which leads to the formation of bleb after cytoplasm flowing to gap between the cell membrane and the cortex. The results showed that the damping coefficient of cortex plays a significant role on the bleb formation. To simulate the experimental phenomena (11), the fracture angle is set to be 90° to fit the width of the pseudopod, the damping coefficient can be selected from a range between 10 pN·s/μm^3^ and 300 pN·s/μm^3^ to get a suitable length of the pseudopod. The generation of bleb is related to cytoplasmic rheology, mechanical properties of the cortex (49), and the tension of the membranes.

As shown in Fig. 19A and Fig. 19B, the tension of the leading edge of the cell membrane begins to decrease with the increase of the local damping coefficient, and the tension change of the trailing edge of the cell remains basically unchanged. That is, with the increase of the local damping coefficient, the leading edge of the cell will have a large deformation state, while the trailing edge will have basically no deformation. The local damping coefficient of the anterior edge of the cortical membrane changes and decreases, but the effect is not large, which can cause the cortex to exhibit a certain contraction; the increase in the local damping coefficient tends to be consistent with the tension change of the posterior edge of the cortex, such as Fig. 19C and Fig. 19D.

**Figure 19.**
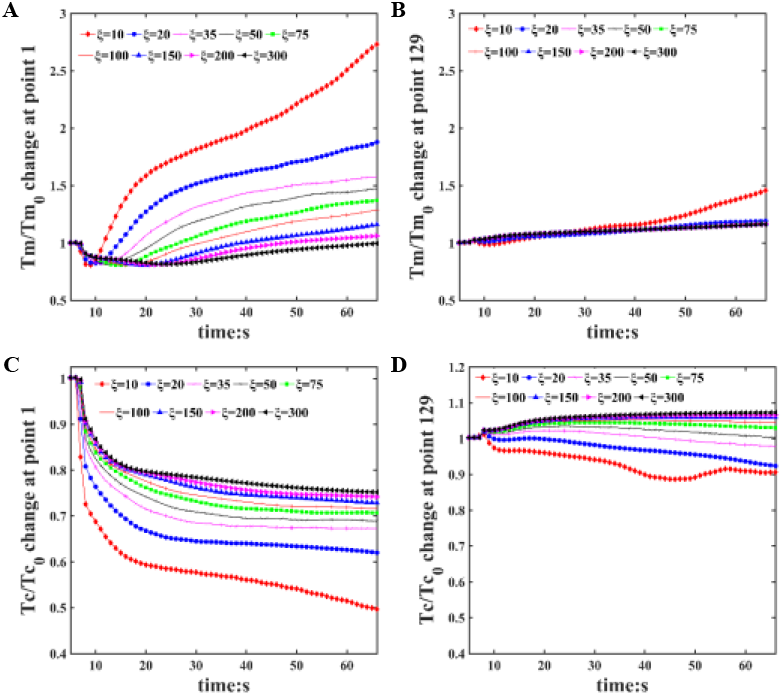
Changes in cell membrane and cortex tension. (A, B) Under different local damping coefficients, the change of cell membrane tension at 1^st^ point on the leading edge and 129^th^ point on the trailing edge, respectively; (C, D) Under different local damping coefficients, the change of cortex membrane tension at 1^st^ point on the leading edge and 129^th^ point on the trailing edge, respectively.

However, when the local damping coefficient is 10 pN·s/μm^3^, the tension of the leading edge of the cell membrane and the cortical membrane changes significantly, and the tension of the trailing edge of the cell membrane also begins to change significantly over time. The tension at the trailing edge of the cortex increases with the increase of the local damping coefficient. However, when the local damping coefficient is less than 35 pN·s/μm^3^, the cortex is in a contracted state. When the local damping coefficient is greater than 35 pN·s/μm^3^, the cortex tension increases. However, the overall variation of the posterior edge of the cortex is between 0.9 ~ 1.1, which has little effect on the overall cell deformation.

## 7. CONCLUSION

In the present paper a two-dimensional fluid-solid coupling cell model is proposed, a numerical solution method for the proposed model basing on immersion boundary numerical method is developed. With the developed numerical method, the formation and evolution of cell bleb caused by broken bonds between cell membrane and cortex are simulated. The number of broken bonds, cortex viscosity coefficient, cell membrane modulus and other parameters that affect bleb production are explored numerically. The effects of these parameters on the size and shape of cellular pseudopod were also investigated. The study concluded the following:

1. With given parameters which are not easily measured experimentally, the experimental bleb of a cell both in size and shape is well simulated, which means that the two-dimensional fluid-solid coupled cell model and numerical solving method developed in this paper are effective for the simulation of cell bleb. The intrinsic mechanics characters are revealed during bleb formation and evolution by considering factors such as cell fluid viscosity, cortex damping coefficient, cell membrane elastic modulus and cortex elastic modulus by the model;
2. The number of broken bonds and the damping coefficient of the cortex have important effects on the bleb size and growth speed. The damping coefficient influence both the fluid and solid in the model, and the fracture angle mainly affects the solid. The pressure and velocity of the fluid in the cell and the length and velocity of the bleb extension increase as the local damping coefficient of the cortex decreases. The tension of the leading edge of the cell membrane increases with the decrease of the fracture angle, but the tension fluctuation of the leading edge of the cortex increases.

## AUTHOR CONTRIBUTIONS

J.J.F. performed the coding, simulations and wrote the article. L.Q.T. supervised and computational model. Z.J.L. supervised and coded. S.B.D. directed the design research. L.C.Z., Y.P.L. and Z.Y.J. contributed to the computational model and the analysis.

## ACKNOWLEDGENTS

The study is financially supported by the National Natural Science Foundation of China (Grant Nos. 11932007, 11772132, 11772131, 11772134, 11972162, and 11602087), and the Science and Technology Program of Guangzhou, China (Grant No. 201903010046).

